# Production of alkanes from CO2 by engineered bacteria

**DOI:** 10.1101/346536

**Authors:** Tapio Lehtinen, Henri Virtanen, Suvi Santala, Ville Santala

## Abstract

**Background:** Microbial biosynthesis of alkanes is considered a promising method for the sustainable production of drop-in fuels and chemicals. Carbon dioxide would be an ideal carbon source for these production systems, but efficient production of long carbon chains from CO_2_ is difficult to achieve in a single organism. A potential solution is to employ acetogenic bacteria for the reduction of CO_2_ to acetate, and engineer a second organism to convert the acetate into long-chain hydrocarbons.

**Results:** In this study, we demonstrate alkane production from CO_2_ by a system combining the acetogen *Acetobacterium woodii* and a non-native alkane producer *Acinetobacter baylyi* ADP1 engineered for alkane production. Nine synthetic two-step alkane biosynthesis pathways consisting of different aldehyde- and alkane-producing enzymes were combinatorically constructed and expressed in *A. baylyi.* The aldehyde-producing enzymes studied were AAR from *Synechococcus elongatus,* Acr1 from *A. baylyi,* and Ramo, a putative dehydrogenase, from *Nevskia ramosa.* The alkane-producing enzymes were ADOs from *S. elongatus* and *Nostoc punctiforme,* and CER1 from *Arabidopsis thaliana.* The performance of the pathways was evaluated with a twin-layer biosensor, which allowed the monitoring of both the intermediate, fatty aldehyde, as well as the alkane production. The highest alkane production, as indicated by the biosensor, was achieved with a pathway consisting of AAR and ADO from *S. elongatus.* The performance of this pathway was further improved by balancing the relative expression levels of the enzymes in order to limit the accumulation of the intermediate fatty aldehyde. Finally, the acetogen *A. woodii* was used to produce acetate from CO_2_ and H_2_, and the acetate was used for alkane production by the engineered *A. baylyi,* thereby leading to the net production of long-chain alkanes from CO_2_.

**Conclusions:** A modular system for the production of drop-in liquid fuels from CO_2_ was demonstrated. Among the studied synthetic pathways, the combination of ADO and AAR from *S. elongatus* was found to be the most efficient in heterologous alkane production in *A. baylyi.* Furthermore, limiting the accumulation of the fatty aldehyde intermediate was found to be beneficial for the alkane production.

## Background

Microbially produced alkanes have been proposed as potential replacements for the current fossil derived fuels and chemicals. Alkanes would be ideal substitutes for currently used fossil fuels, because they can drop in directly in the current infrastructure and engines. However, realization of alkane production in environmentally viable way requires that alkanes are produced from sustainable substrates with high efficiency. Currently, most of the microbial alkane production studies have focused on alkane production from biomass-derived sugars. In contrast, carbon dioxide could allow to bypass the biomass step and serve as sustainable carbon source for large-scale production, provided that efficient methods and microbial hosts can be developed for its utilization.

The challenge of alkane production from carbon dioxide can be divided in two parts: 1) the reduction of carbon dioxide to organic compounds, and 2) the conversion of these compounds to long-chain alkanes. These processes impose different, and in part contrasting, requirements for the host metabolism, and are thus regarded difficult to achieve efficiently in any single organism. For example, the most efficient CO_2_ fixation pathways require anaerobic conditions and elevated CO_2_ concentrations [1,2]. Due to the anaerobic metabolism, the organisms employing these pathways have limited capacity for the production of highly energy-intensive molecules, such as long-chain alkanes [3].

Acetogenic bacteria (acetogens) are known for their ability to efficiently reduce carbon dioxide to acetate by the Wood-Ljungdahl pathway [4]. The energy for the acetogenesis can be obtained from hydrogen gas, organic compounds, or in the form of electrons directly from the cathode surface of a bioelectrochemical system in a process called microbial electrosynthesis [5,6]. Acetogens have been studied for decades and are already being used in industrial scale [7]. Thus, acetogens present an attractive option for the fixation of carbon dioxide into organic compounds. However, due to the limitations of their anaerobic energy metabolism, they are unlikely to be able to efficiently produce molecules with higher energy content [8,9].

Furthermore, metabolic engineering of acetogens is hampered by the lack of established gene editing tools. One solution to circumvent these issues and to broaden the metabolic capability is to employ more than one organism in the production system. For example, the acetate produced from CO_2_ by the acetogens can be upgraded to value-added, high-energy molecules by heterotrophic organisms such as *Acinetobacter baylyi* [10] or *Yarrowia lipolytica* [11]. Hu *et al.* produced acetate with *Moorella thermoacetica* from CO_2_ and H_2_, and utilized *Y. lipolytica* for the conversion of the acetate into triacylglycerol. We have previously studied the conversion of acetate produced by acetogens either from CO_2_ and H_2_, or by CO_2_ and electricity (microbial electrosynthesis) into long alkyl chains. However, alkane production from acetate has not been reported to date.

Alkanes have been produced in various microbial hosts by the heterologous expression of a cyanobacterial two-step pathway. The pathway involves the reduction of fatty acyl-ACP to fatty aldehyde, which is subsequently converted to n-alkane. In cyanobacteria, the first step is catalyzed by acyl-ACP reductase (AAR) and the second step by aldehyde deformylating oxygenase (ADO) [12]. Besides the cyanobacterial AAR homologs, fatty aldehyde-producing reductases have been characterized from various organisms [13–15]. Even though these reductases are not known to participate in alkane production in their native context, they can be utilized as part of synthetic alkane producing pathways in heterologous hosts. Similarly, an alkane-producing enzyme CER1 from *Arabidopsis thaliana* has been used in heterologous alkane production instead of the cyanobacterial ADOs [16,17].

The last enzyme of the pathway, ADO, is regarded as the bottleneck for the alkane biosynthesis, due to its low activity [18]. Cyanobacteria naturally produce only small amounts of alkanes, and thus the low activity is not an issue in the native context. However, in the (heterologous) overproduction, the low activity of ADO can cause accumulation of the pathway intermediates. Especially the fatty aldehyde, which is the substrate for ADO, is problematic due to its toxic nature [19]. In addition, endogenous enzymes, such as aldehyde reductases [20] and dehydrogenases [21], compete for the fatty aldehyde substrate with ADO, leading to the loss of the valuable long acyl chains from the alkane-producing pathway. Thus, efficient alkane production requires careful balancing of the enzyme activities in order to provide ADO with sufficient amount of substrate, while avoiding these negative effects of fatty aldehyde accumulation. Furthermore, while the individual components (especially the cyanobacterial AARs and ADOs) of the pathway have been thoroughly studied [22–24], less attention has been paid to the performance of synthetic pathways comprised of enzymes from different origins.

We have previously developed a twin layer biosensor for the monitoring of the alkane biosynthesis pathway [25]. The sensor consists of two parts, and is able to monitor both the intermediate fatty aldehyde and the end product alkane. The first part of the sensor consists of constantly expressed bacterial luciferase LuxAB, which utilizes fatty aldehyde as a substrate in a reaction that produces visible light. Thus, the concentration of fatty aldehydes is reflected in the luminescence signal. The second part of the sensor consists of a *gfp* gene under a native alkane-inducible promoter. Both luminescence and fluorescence can be measured in real time during cell growth, making the sensor a powerful tool for the screening and optimization of the alkane biosynthesis pathways in a high-throughput manner.

*A. baylyi* ADP1 is a promising chassis for genetic studies, metabolic engineering, and lipid production [26–30]. The strain naturally produces storage lipids, and thus the central metabolism is wired for the efficient production of fatty acyl-derived molecules. *A. baylyi* does not naturally produce alkanes, but the fatty aldehyde intermediate is part of its natural storage lipid metabolism [31]. We have previously shown that alkanes can be produced by the expression of the cyanobacterial pathway (AAR and ADO) [25]. In addition, A. baylyi grows well on acetate, and we have previously shown it to be a suitable chassis for the upgrading of the acetate produced by acetogens into long-chain lipid compounds [10]. Furthermore, *A. baylyi* ADP1 has tendency for natural transformation and homologous recombination, supporting straightforward genome editing [26].

In this study, we establish alkane production from acetate in metabolically engineered *A. baylyi* ADP1. By utilizing the alkane biosensor, we compare the performance of different synthetic alkane biosynthesis pathways, and demonstrate the importance of pathway balancing for improved alkane production. Further, we demonstrate the net production of alkanes from CO_2_ and H_2_ by a two-stage bacterial process combining an acetogen *Acetobacterium woodii* and the *A. baylyi* engineered for alkane production (Figure 1).

**Figure 1.**
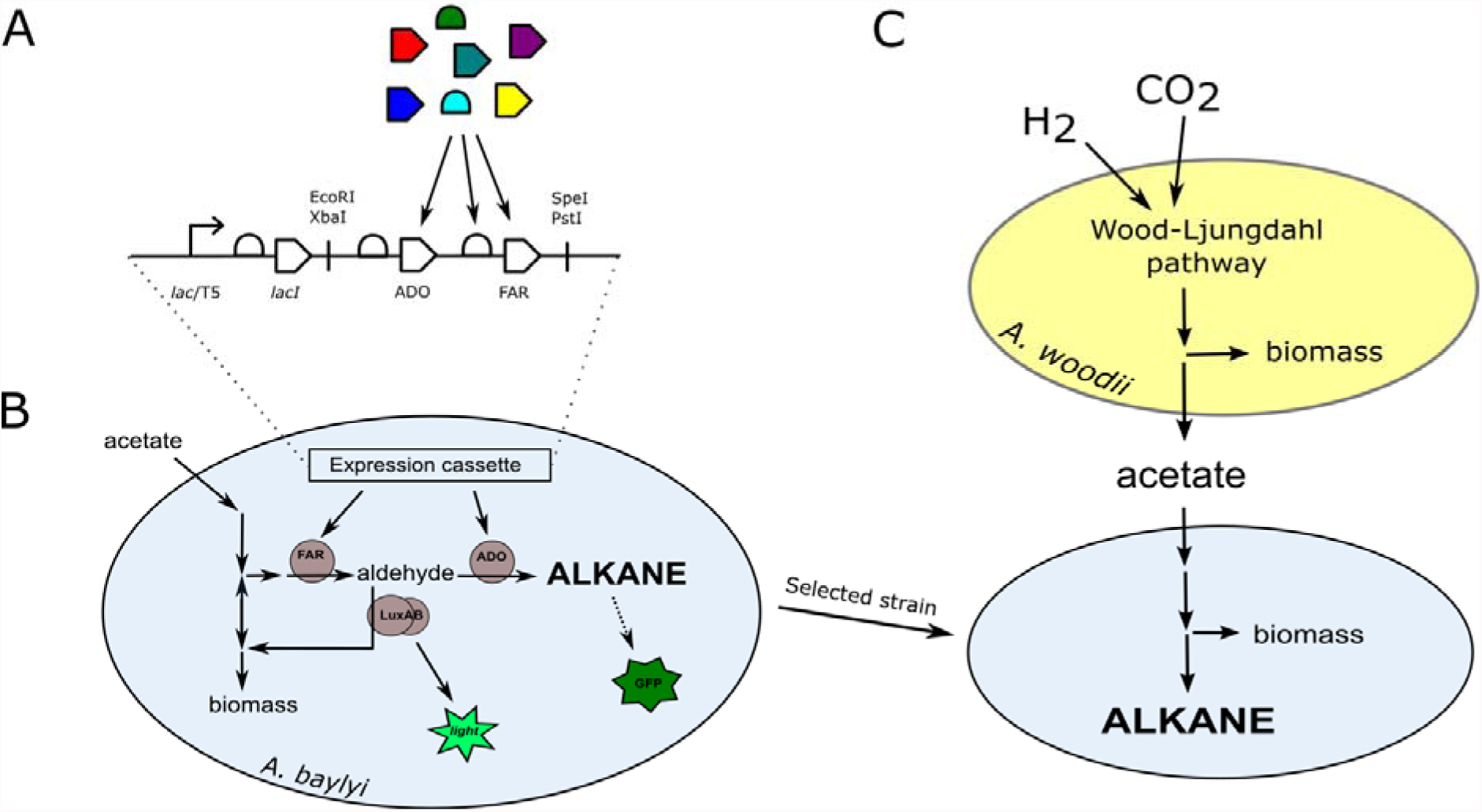
Outline of the study. Nine synthetic alkane-producing pathways were constructed from genetic parts originating from different species (A). The performance of these pathways in alkane production from acetate was evaluated by a previously developed twin-layer sensor (B), which allows the detection of both the pathway intermediate (fatty aldehyde) and the end product (alkane). The engineered strain that produced most alkanes was coupled with a process in which acetate was produced from carbon dioxide by acetogenic bacteria, leading to the net production of alkanes from carbon dioxide (C).

## Results

As the first step towards the alkane production from CO_2_, we started by constructing nine synthetic alkane biosynthesis pathways and compared their performance in alkane production from acetate. We have previously demonstrated increased fatty aldehyde production in *A. baylyi* with the expression of any of the three different reductases: Acr1, AAR and Ramo [32]. Acr1 is an endogenous fatty acy1-CoA reductase from *A. baylyi* that is involved in the natural storage lipid (wax ester) production pathway. AAR from *Synechococcus elongatus* is a fatty acyl-ACP/CoA reductase employed in the alkane production pathway in cyanobacteria, and earlier studies indicate that it is the most efficient among the cyanobacterial homologs [22]. Ramo, on the other hand, is a putative short-chain dehydrogenase from the bacterium *Nevskia ramosa.* Ramo was initially identified based on its sequence homology with Acr1, and its expression was shown to increase the fatty aldehyde and wax ester production in *A. baylyi* [32]. These three fatty aldehyde-producing enzymes were included in this study. The alkane-producing enzymes selected for the study were ADOs from *Synechococcus elongatus* (SeADO) and *Prochlorococcus marinus* (PmADO), and CER1 from *Arapidopsis thailana,* all of which have been previously utilized in heterologous alkane production.

The genes encoding the pathway enzymes were cloned under T5*/lac* promoter as synthetic operons containing first the alkane-producing and then the aldehyde-producing gene, both preceded by ribosomal binding site (RBS) sequences (Fig 1b). The resulting nine constructs were transformed to the previously developed biosensor strain [25]. *A. baylyi* naturally degrades alkanes, but the *alkM* gene essential for alkane degradation [33] has been deleted from the sensor strain [25].

For the comparison of alkane production, the sensor cells harboring the different pathways were cultivated in 96-well plate in minimal medium supplemented with 50 mM acetate. The expression of the alkane biosynthesis genes was induced with either 10 or 100 μM IPTG and the luminescence and fluorescence signals measured in real time during cell growth (Fig 2). The highest luminescence signal, indicating highest fatty aldehyde production, was obtained with the CER1-AAR pair. On the contrary, the highest fluorescence signal, indicating the highest alkane production, was obtained with SeADO-AAR pair with the 100 μM IPTG induction. This pair was selected for further studies.

**Figure 2.**
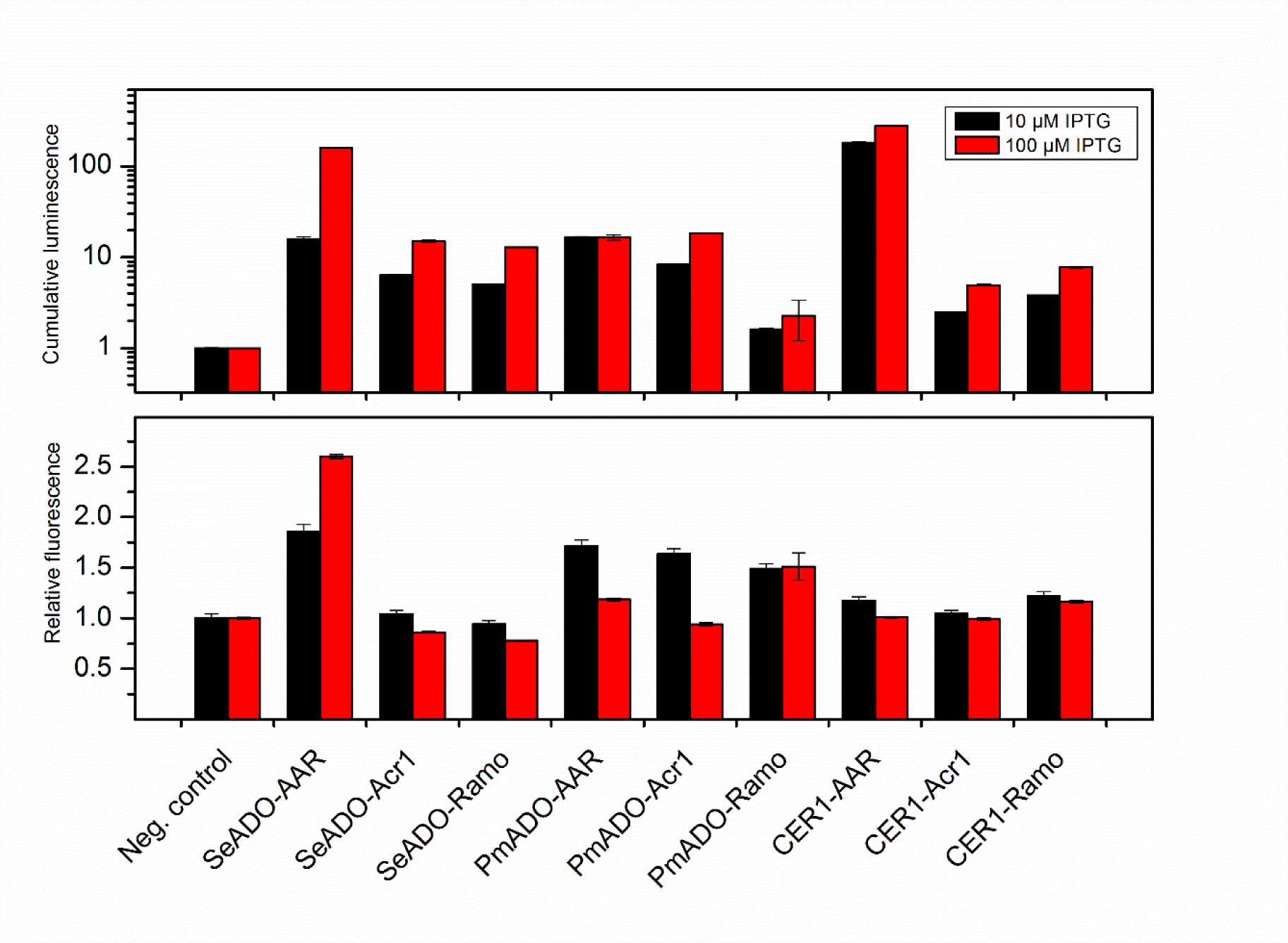
Comparison of the synthetic pathways. The sensor cells containing different alkane biosynthesis pathways were cultivated in minimal medium with 50 mM acetate. The expression of the alkane biosynthesis genes were induced with 10 or 100 μM IPTG, and the luminescence (upper panel) and fluorescence (lower panel) signals produced by the sensor cells were measured. The cumulative luminescence signal reflects the amount of fatty aldehyde produced by the cells throughout the cultivation (relative to the control), and the relative fluorescence signal reflects the activation of the *gfp* expression in response to the alkanes produced by the cells. The negative control strain is the sensor strain without the alkane biosynthesis pathway. Average and standard deviation of two replicates are shown.

Optimal production requires that the enzyme activities in the pathway are balanced, thus avoiding the accumulation of intermediates while maintaining high flux through the pathway. Thus, we next set out to optimize the relative expression levels of ADO and AAR. Changes in the relative expression was achieved by modulating the RBS of AAR in the synthetic operon. Three RBS:s with predicted strengths “weak”, “medium” and “strong” based on *E. coli* characterization were chosen (Biobrick parts BBa_0031, BBa_0032, and BBa_0030, respectively). The original strong RBS (BBa_0034) of ADO was not changed. The three synthetic operons with varying AAR RBSs were again expressed in the sensor strain. The cells were cultivated in the 96-well plate, and luminescence and fluorescence measured in real time. As expected, the highest luminescence signal was obtained with the strongest RBS, indicating that the highest expression level of AAR led to the highest fatty aldehyde production (Fig 3, upper panel). In contrast, the highest fluorescence signal was obtained with the intermediate strength RBS, suggesting that relatively lower expression of AAR leads to more balanced metabolism and higher alkane production (Fig 3, lower panel). Thus, this strain was utilized in the subsequent alkane production experiments.

**Figure 3.**
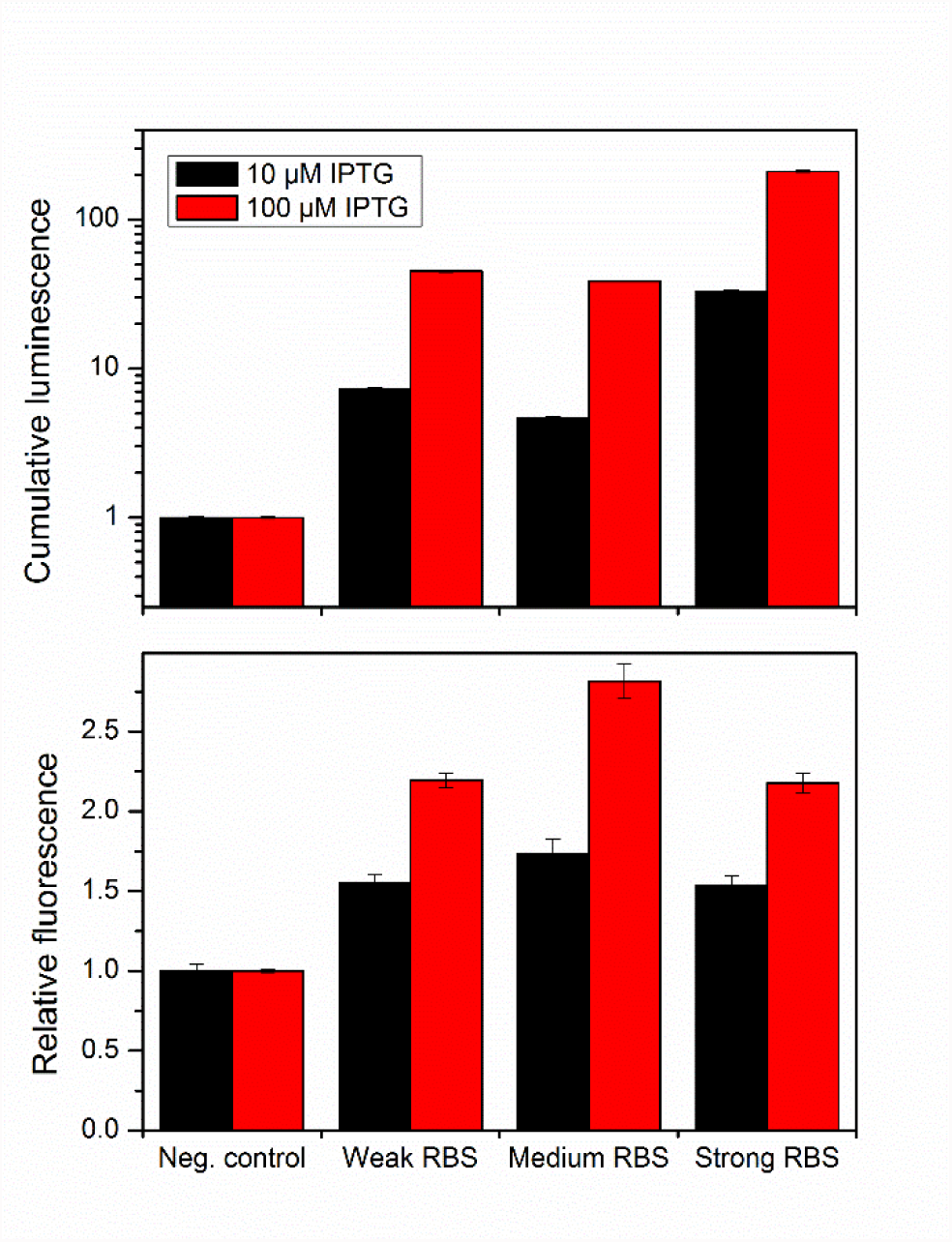
Balancing the relative expression levels of ADO and AAR. The alkane biosynthesis pathway consisting of SeADO and AAR was balanced by altering the ribosomal binding site (RBS) of AAR, while the RBS of SeADO was remained unchanged (strong). The sensor cells harboring the pathways with different RBS strengths were cultivated in minimal medium with 50 mM acetate. The expression of the alkane biosynthesis genes were induced with 10 or 100 μM IPTG, and the luminescence (upper panel) and fluorescence (lower panel) signals were measured. The cumulative luminescence signal reflects the amount of fatty aldehyde produced by the cells throughout the cultivation (relative to the control), and the relative fluorescence signal reflects the activation of the *gfp* expression in response to the alkanes produced by the cells. The negative control strain is the sensor strain without the alkane biosynthesis pathway. Average and standard deviation of two replicates are shown.

Acetate is a challenging substrate for many microbes due to its toxic nature [34]. Although *A. baylyi* grows well on reasonably high acetate concentrations [10,35], carbon limitation is still inevitable in batch cultivations. In order to circumvent this issue and allow higher growth and alkane production, we next established a fed-batch cultivation strategy in a 1-liter benchtop bioreactor. The engineered alkane production strain was cultivated in the reactor in minimal medium. The cultivation was initiated with a batch-phase in 500 ml of the medium with 0.2% casein amino acids as the sole carbon source. After that, the expression of the alkane biosynthesis operon was induced with 100 μM IPTG and the feed initiated. The feed solution contained 80 g/L acetic acid, and the rate was manually controlled to keep the acetate concentration in the reactor between 30 and 60 mM in order to avoid carbon limitation. The cultivation was continued for 25 hours after the beginning of the feed, and cell growth, acetate consumption, and alkane production were monitored (Fig. 4). During the cultivation, the cells consumed approximately 400 mM acetate (Fig 4). The final heptadecane titer was 540 μg/L (productivity 21 μg/(L*h)).

**Figure 4.**
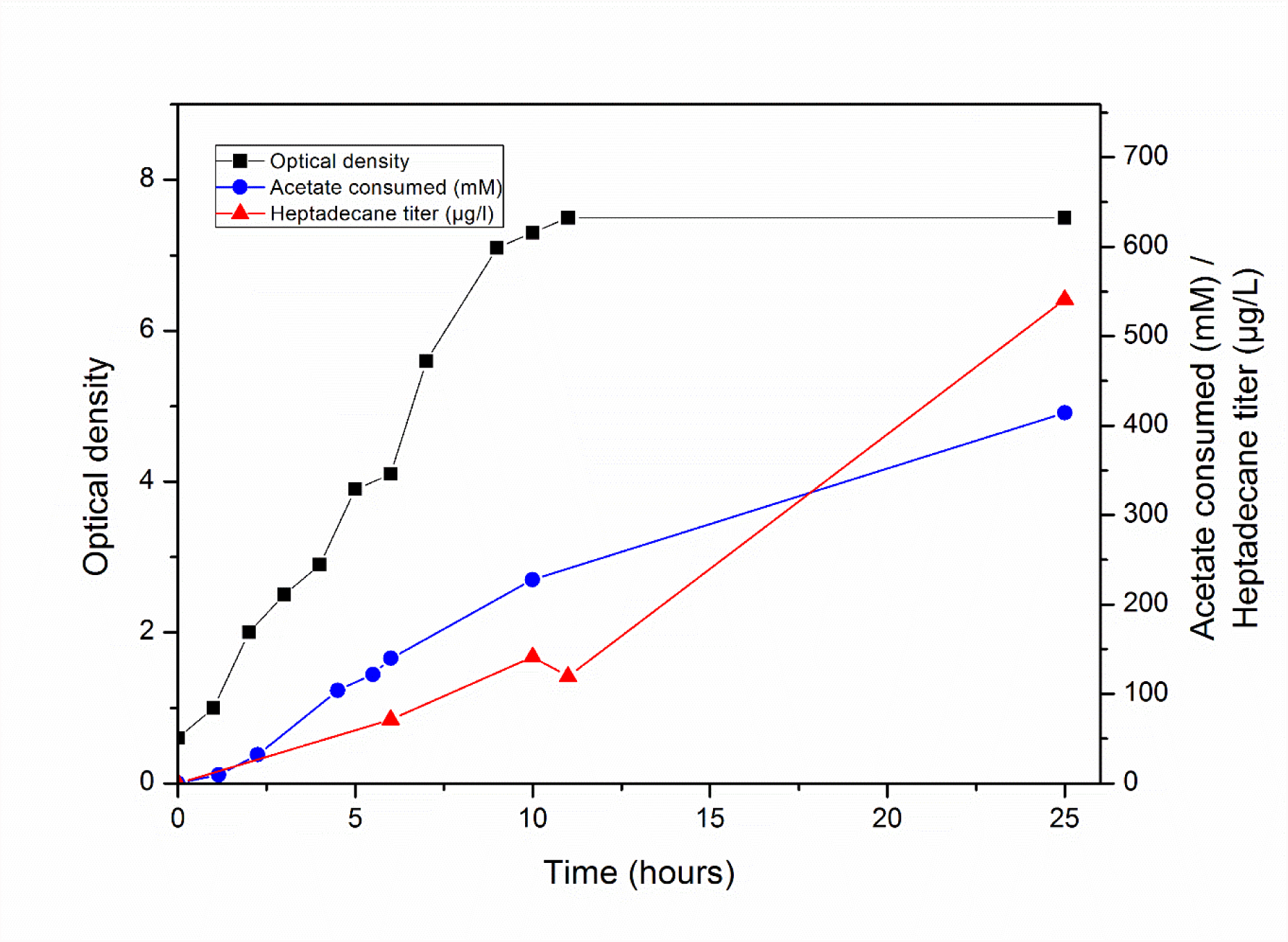
Alkane production from acetate. The selected alkane-producing strain was cultivated in bioreactor in minimal salts medium in fed-batch mode. The feed contained 80 g/L acetic acid, and the feed rate was manually controlled to keep the acetate concentration in the reactor between 30 and 60 mM. Acetate consumption was calculated based on the measured acetate concentrations in the reactor and the volume of the acetic acid solution fed to the reactor.

Acetogens are known for their ability to efficiently reduce carbon dioxide to acetate. In order to achieve alkane production from carbon dioxide, we employed the acetogen *A. woodii* for the production of acetate from CO_2_ and H_2_. *A. woodii* was grown in a bioreactor in 600 ml of minimal medium, and the medium was continuously flushed with H_2_/CO_2_ gas. During a six-day cultivation, *A. woodii* grew to an optical density of 2.3 and produced approximately 190 mM acetate (Fig 5a). The medium from this cultivation, including the *A. woodii* biomass, was used as the feed for the alkane-producing A. *baylyi* as such. *A. baylyi* was grown in a bioreactor similarly as earlier. After the initial batch phase with 0.2% casein amino acids as the carbon source, the feed was initiated, and the alkane production induced with 100 μM IPTG. The rate of the feed was manually controlled to keep the acetate concentration in the reactor between 30 and 70 mM. Due to the relatively dilute feed, the volume of the cultivation increased from an initial 300 ml to the final 800 ml. *A. baylyi* utilized the feed for growth and alkane production (Fig 5b), resulting in the net production of alkanes from CO_2_ and H_2_. During the 9-hour cultivation, *A. baylyi* produced 74 μg/L heptadecane (productivity 8.2 μg/(L*h)), and consumed approximately 92 mM of acetate.

**Figure 5.**
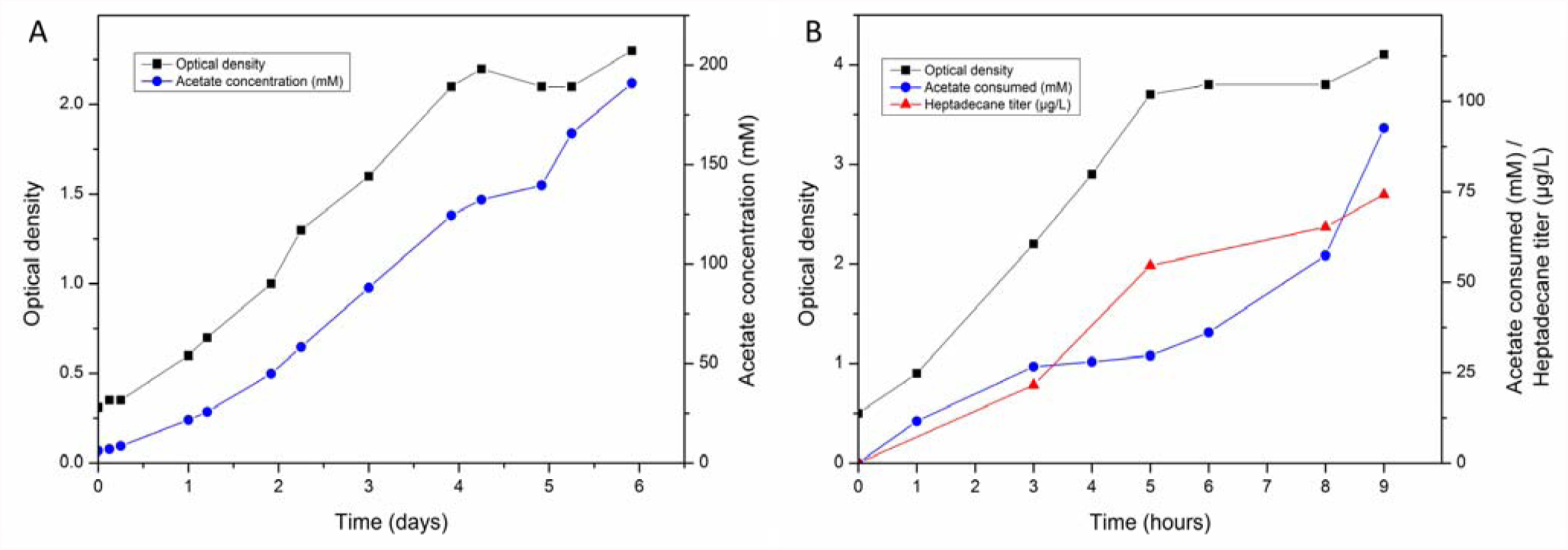
Alkane production from CO_2_ and H_2_. *Acetobacterium woodii* was grown in minimal medium with CO_2_ and H_2_, and cell growth and acetate production measured (A). After six days, the acetate-containing *A. woodii* medium was used as the feed for alkane-producing *A. baylyi* in fed-batch culture (B). The feed rate was manually controlled to keep the acetate concentration in the medium between 30 and 70 mM. *A. baylyi* utilized the feed for growth and alkane production (B). Due to the relatively dilute feed (containing approximately 190 mM acetate), the culture volume of the *A. baylyi* cultivation increased from 300 ml to 800 ml during the process.

## Discussion

In the production of bulk chemicals, such as liquid fuels, required in massive quantities, the sustainability of the raw material is of special importance. Carbon dioxide would be an ideal carbon source due to its abundancy, independency of land usage, and the urgent need to limit its concentration in the atmosphere. On the other hand, alkanes could serve as drop-in replacements to current, fossil derived, liquid fuels. Microbes hold potential for both the production of alkanes, and the reduction of CO_2_, but significant advancements are required in order to establish an efficient microbial process for the conversion of CO_2_ to alkanes.

In this study, we demonstrated the microbial production of alkanes from CO_2_ and H_2_ by a system combining the strengths of two different bacterial species. For the CO_2_ reduction, we employed the acetogen *A. woodii,* which is able to efficiently produce acetate from CO_2_ and H_2_ [36]. To convert the acetate to alkanes, and to achieve the net production of alkanes from CO_2_, we studied heterologous alkane production in *A. baylyi* ADP1, which has been previously used to upgrade acetate to long-chain lipid molecules [10].

Microbial alkane production can be achieved in heterologous hosts by the expression of aldehyde-producing fatty acyl-CoA/ACP reductase (AAR) and aldehyde deformylating oxygenase (ADO) [37]. We have previously found that the fatty aldehydes produced in *A. baylyi* may have different metabolic fates depending on the reductase that is producing them. Specifically, the aldehydes produced by Acr1 or Ramo were efficiently directed to fatty alcohols and further to wax esters by native *A. baylyi* enzymes, but the aldehydes produced by cyanobacterial AAR were not [32]. Furthermore, it has been speculated previously that the cyanobacterial AAR and ADO might interact with each other and thus ensure efficient delivery of the fatty aldehyde from AAR to ADO [24]. Thus, it is reasonable to assume that there might be differences in the efficiency of the aldehyde delivery from AAR to ADO in synthetic pathways utilizing enzymes originating from different species. In order to investigate the function of alkane biosynthesis pathways consisting of different enzymes, we started by constructing and comparing nine synthetic pathways that comprised of different aldehyde- and alkane-producing enzymes.

For the comparison of the synthetic pathways, we utilized the previously developed twin-layer alkane biosensor, which allows the monitoring of both the aldehyde intermediate and the end product alkane. The monitoring of the aldehyde intermediate, in addition to the end product alkane, yields valuable information about the functionality of the aldehyde-producing enzymes in this context, and enables rational evaluation of potential pathway bottlenecks. Three aldehyde-producing reductases (AAR, Acr1, Ramo) and three alkane-producing enzymes (SeADO, PmADO, CER1) were selected for the study. The expression of any of the aldehyde-producing reductases in combination with any of the alkane-producing enzymes was associated with aldehyde production, as measured by the biosensor. The results are in agreement with our earlier study, where the fatty aldehyde production was increased in *A. baylyi* by the overexpression of any of these reductases [32].

Based on the biosensor output, there are significant differences on the efficiency of alkane production from the aldehydes depending on the enzyme pair. The combination of CER1-AAR led to the highest aldehyde production, whereas the SeADO-AAR pair led to the highest alkane production. Interestingly, alkane production by SeADO was significantly higher when AAR was used to provide the aldehydes, than with the other two reductases (Acr1 or Ramo). With the other two alkane-producing enzymes (PmADO, and CER1), the difference was not so clear, and they seem to accept the aldehydes equally well from any of the reductases tested.

For synthetic pathways to work efficiently, the enzyme activities need to be balanced in order to avoid the accumulation of pathway intermediates and the metabolic burden caused by unnecessary high protein expression [38]. In the case of the alkane biosynthesis pathway, the relatively low activity of ADO compared to AAR may lead to the accumulation of the fatty aldehyde, potentially hampering the efficiency of alkane production. Thus, we next set out to balance the relative expression of ADO and AAR for improved alkane production. The relative expression of AAR was altered by modifying the strength of the RBS, and the effects on the aldehyde and alkane production were measured with the biosensor. The strongest RBS yielded the highest aldehyde production, as expected. In contrast, the alkane production was higher when an RBS with intermediate strength was used. This suggests that lowering the AAR expression leads to more balanced metabolism, thus avoiding the harmful effects of the fatty aldehyde accumulation, and resulting in higher alkane production. This result is in agreement with a recent study, where it was concluded that the accumulation of fatty aldehydes was not beneficial for the alkane production in *E. coli* [39].

Alkane biosensors have been suggested as a potential tool for speeding up the strain development and the screening of novel and/or altered enzymes for alkane production [37]. Even though various alkane biosensors have been developed [25,40,41], this is the first time a sensor is used for the evaluation of the performance of different pathway configurations. The twin-layer sensor utilized here allows the monitoring of both the key intermediate and the end product of the alkane biosynthesis pathway, offering insights into the function of the pathway. The successful utilization of the sensor for the pathway balancing exemplifies the benefits of biosensors in the development of efficient production strains.

In many microbes, including *A. baylyi,* efficient lipid production is promoted by an excess of the carbon source. Typically, favorable conditions are achieved by maintaining a high carbon-to-nitrogen ratio in the culture medium. With acetate, however, high concentrations are not feasible due to its toxicity. Consequently, batch cultivations are rapidly depleted of acetate, leading to carbon limitation and inefficient lipid production. In this study, we utilized a fed-batch cultivation strategy in order to support better growth and alkane production from acetate. In the cultivations, acetate concentration was maintained between 30 and 70 mM in order to provide the cells with sufficient carbon and energy source while avoiding the toxic effects of higher concentrations. With this strategy, alkanes could be produced both from pure acetic acid and from the acetic acid produced by the acetogen *A. woodii.* In our experiments, the alkane production was more efficient when pure acetic acid was used as the feed when compared to the medium from the acetogenic cultivation. One explanation is that the used medium also contained the biomass of *A. woodii,* leading to relatively lower carbon-to-nitrogen ratio, potentially negatively effecting the alkane production.

We have previously studied wax ester (WE) production from acetate in *A. baylyi* [10]. WEs are produced from the same intermediates that are used in the alkane production pathway (fatty aldehyde and fatty acyl-CoA), and thus it is interesting to compare the efficiency of WE and alkane production. The WE titer from acetate (ca. 90 mg/L, [10]) was significantly higher than the alkane titer in the present study (< 1 mg/L), suggesting that the precursor supply is not limiting the alkane production even in the relatively low-carbon conditions. This implies that future research should focus on improving the last step (conversion of fatty aldehydes to alkanes) in the alkane production pathway. For example, the activity of ADO may be improved by the coexpression of an electron transfer system [42].

Acetate is a platform chemical that can be produced from CO_2_ by various microbial processes, such as microbial electrosynthesis or syngas fermentation. Throughout this study we utilized acetate as the carbon source for alkane production, in order to allow the connection of the alkane-producing module to an upstream acetate-producing module. As a proof of principle, in the final stage of this study, we connected the alkane production with the upstream acetate production from CO_2_ by the acetogen *A. woodii.* The two-stage production system converges around acetate, and enables the construction of versatile processes by plugging different acetate-producing and -consuming modules in the system. For example, the final product can be modified by engineering the production host. We have previously studied the wax ester production from acetate produced by the acetogens [10], while other studies have reported the production of triacylglycerides [11], or n-butanol, polyhydroxybutyrate, and isoprenoids [43]. In this study, aliphatic long chain alkane production from acetate was demonstrated, further expanding the product range to hydrocarbons, enabling the production of drop-in replacements for liquid fuels. The modular nature of the two-stage production system brings the additional advantage that both the CO_2_ fixation and alkane production can be optimized independently of each other. For example, significantly higher acetate production rates have been obtained with *A. woodii* with optimized reactor design [36] or metabolic engineering [44]. In this work, we focused on the alkane production from acetate by *A. baylyi,* but the overall productivity and titer of the system could be enhanced by optimizing both modules. As an easy-to-engineer chassis, *A. baylyi* can potentially be utilized for the production of a range of compounds, further emphasizing the modular nature of the system.

## Conclusions

In this study, we demonstrated the production of long-chain alkanes from CO_2_ by a bacterial system comprising of two modules. In the first module, CO_2_ is reduced to organic acids by acetogenic bacteria. In the second module, these organic acids are converted to long-chain hydrocarbons by the A. *baylyi* engineered for alkane production. The modular nature of the system allows both the acetate-producing and -consuming modules to be improved separately. In this study, we focused on the alkane production by A. *baylyi,* and compared the performance of different synthetic alkane-producing pathways. The utilization of a twin-layer biosensor enabled the balancing of the pathway in order to avoid the accumulation of the intermediate fatty aldehyde, which led to improved alkane production.

## Methods

### Bacterial strains and genetic modifications

The previously described alkane biosensor (ASI) constructed in *Acinetobacter baylyi* ADP1 (DSM 24139) [25] was used for the construction of alkane-producing strains. For molecular cloning and plasmid maintenance, *Escherichia coli* XL1-Blue (Stratagene, USA) was used. *Acetobacterium woodii* (DSM 1030) was ordered from Deutsche Sammlung von Mikroorganismen und Zellkulturen (DSMZ).

For the expression of the alkane biosynthetic genes, a previously described expression cassette was used [32]. The cassette contains an IPTG-inducible T5/*lac* promoter, followed by the *lacl* gene and the BioBrick RFC10-accepting cloning site (EcoRl-Xbal-Spel-Pstl) (Fig 1b). The cassette also contains a spectinomycin resistance marker gene under its own promoter, and flanking sites to facilitate the integration of the cassette to the prophage region of *A. baylyi* [45]. *Acr1* (ACIAD3383) was amplified from the genome of *A. baylyi* ADP1 and *ramo* (an open reading frame encoding for a putative short chain dehydrogenase WP_022976613.1) from the genome of *Nevskia ramosa* (DSM 11499) with primers tl17 & tl18 and tl33 & tl34, respectively (Table 1). A*ar (S. elongatus aar,* WP_011242364.1), *seado* (*S. elongatus ado,* WP_011378104.1), *pmado (P. marinus ado,* WP_011130600.1) *and cer1* (Arabidopsis thaliana, NP_001184890.1) were codon-optimized and purchased from Genscript (USA) with appropriate restriction sites and ribosomal binding sites (Biobrick part BBa_0034). The nucleotide sequences of the genes are provided in the Additional file 1. These ADO and AAR analogs were sequentially cloned to the expression cassette using the Biobrick RFC10 protocol to produce 9 different combinations. For the RBS modulation, the *aar* gene was PCR-amplified with the primers containing the appropriate RBS sequences (Table 1), and cloned to the *seado-*containing expression cassette. The expression cassettes were transformed to the alkane biosensor strain by natural transformation as described previously [29], and colonies were selected with 50 μg/ml spectinomycin. All genetic modifications were confirmed with PCR and sequencing.

**Table 1.**
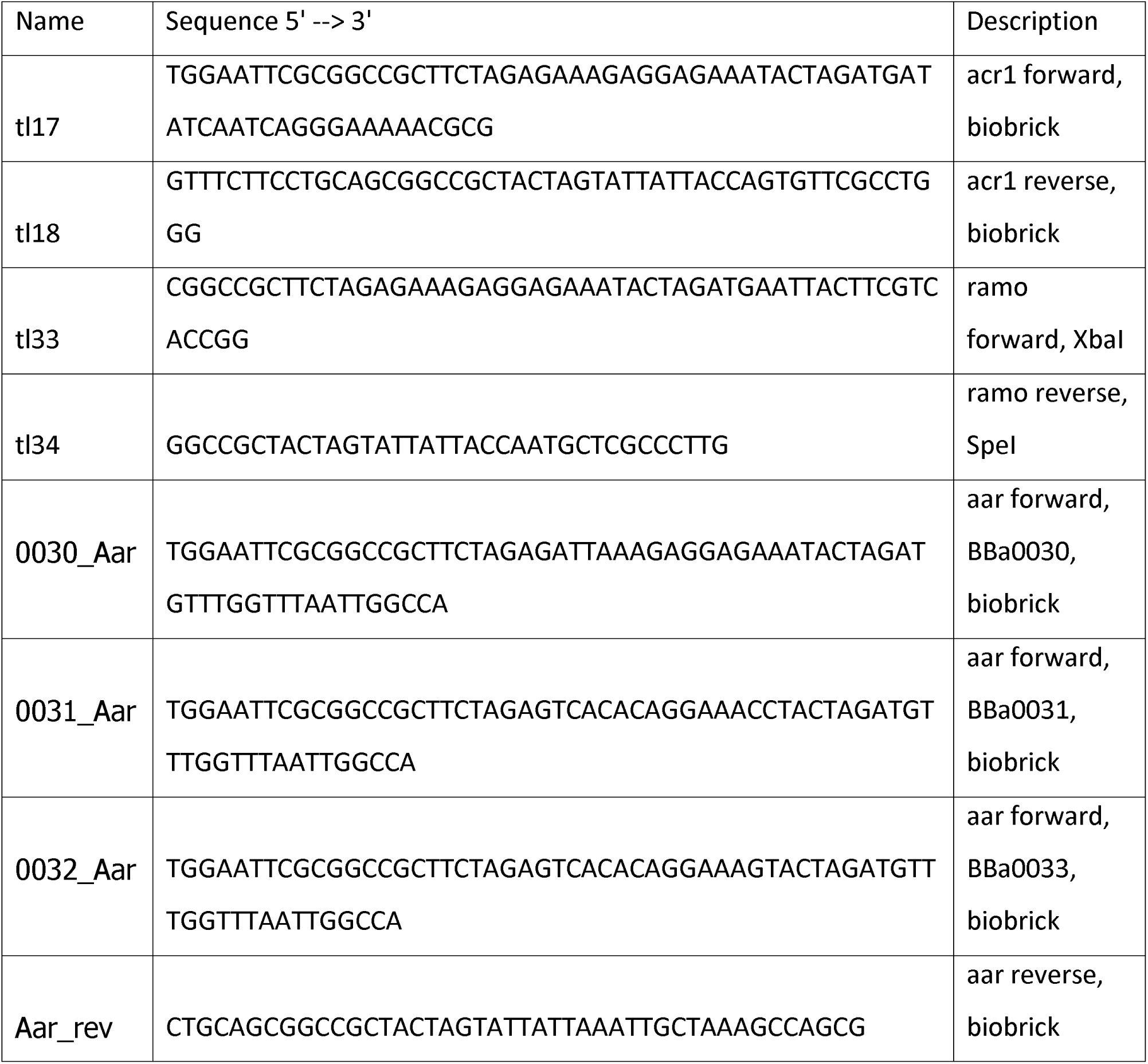
Primers used in the study.

#### Medium and culture conditions

Cloning and routine maintenance of *A. baylyi* and *E. coli* were performed in LB medium (10 g/l tryptone, 5 g/l yeast extract, 5 g/l NaCl). In alkane production experiments, *A. baylyi* was cultivated in modified minimal salts medium MA/9. The medium contained (per liter): 5.52 g Na2HPO4 * 2 H2O, 3.4 g KH2PO4, 1 g NH4Cl, 2 g casein amino acids, 0.008 g nitrilotriacetic acid, 2.0 mg FeCl3, 11.1 mg CaCl2, and 240 mg MgSO4. Acetic acid was used as the carbon source in concentrations indicated in the results section. IPTG (isopropyl β-D-1-thiogalactopyranoside) was used for the induction of the alkane biosynthesis genes in concentrations indicated in the results section. Spectinomycin (50 μg/ml) was used for selection.

For the real time, *in vivo* luminescence and fluorescence measurements, the cells were cultivated in 96-well plate (Greiner Bio-one, Austria) in Spark microplate reader (Tecan, Switzerland). The cells were inoculated from overnight culture to an initial optical density of approximately 0.05 and cultivated in 200 μl of medium. The plate was placed inside a humidity cassette (Tecan) to prevent evaporation. The cultivation and measurement were performed at 30 °C. Optical density (600 nm), luminescence and fluorescence (excitation 485 ± 10 nm, emission 510 ± 5 nm) signals were measured every 30 minutes, and the plate was shaken for 5 minutes twice an hour (108 rpm, 2,5 mm amplitude). The relative fluorescence values were obtained by dividing the fluorescence reading with the optical density. The cumulative luminescence value were calculated by adding all the luminescence readings together.

A. woodii was grown in DSM 135 medium containing (per liter): 1 g NH_4_Cl, 0.33 g KH_2_PO_4_, 0.45 g K_2_HPO_4_, 0.1 g MgSO_4_ × 7 H_2_O, 2 g Yeast extract, 10 g NaHCO_3_, 10 g D-Fructose, 0.5 g L-Cysteine-HCl, 30 mg Nitrilotriacetic acid, 60 mg MgSO_4_ × 7 H_2_O, 10 mg MnSO_4_ × H_2_O, 20 mg NaCl, 2 mg FeSO_4_ × 7 H_2_O, 3.6 mg CoSO_4_ × 7 H_2_O, 2 mg CaCl_2_ × 2 H_2_O, 3.6 mg ZnSO_4_ × 7 H_2_O, 0.2 mg CuSO_4_ × 5 H_2_O, 0.4 mg KAl(SO_4_)2 × 12 H_2_O, 0.2 mg H3BO_3_, 0.2 mg Na_2_MoO_4_ × 2 H_2_O, 0.6 mg NiCl_2_ × 6 H_2_O, 6 μg Na_2_SeO_3_ × 5 H_2_O, 8 μg Na_2_WO_4_ × 2 H_2_O, 20 μg Biotin, 20 μg Folic acid, 100 μg Pyridoxine-HCl, 50 μg Thiamine-HCl × 2 H_2_O, 50 μg Riboflavin, 50 μg Nicotinic acid, 50 μg D-Ca-pantothenate, 1 μg Vitamin B12, 50 μg p-Aminobenzoic acid, and 50 μg Lipoic acid. For autotrophic growth with CO_2_, fructose was omitted, the concentration of yeast extract increased to 4 g/l and NaHCO_3_ decreased to 5 g/l [36].

#### Bioreactor

The bioreactor cultivations were carried out in 1-liter vessel (Sartorius Biostat B plus Twin System, Germany) with a working volume of 600 ml *(A. woodii)* or 300-800 ml *(A. baylyi),* as indicated in the results section. pH was controlled to 7.0 by automated addition of 1 M H_3_PO_4_ or 5 M KOH. Temperature was kept at 30 °C and stirring speed at 300 rpm. For the *A. baylyi* cultivations, air was supplied at 1 l/min, and partial oxygen pressure controlled to 20% saturation by supply of oxygen. For *A. woodii* cultivations, the reactor was operated at anaerobic conditions, and a mixture of H_2_ and CO_2_ (80%/20%) was purged through the system at 1 l/min.

#### Analytical methods

Acetate concentrations were measured with high performance liquid chromatography using LC-20AC prominence liquid chromatograph (Shimadzu, Japan) and Rezex RHM-Monosaccharide H+(8%) column (Phenomenex, USA) with 5mM H_2_SO_4_ as the mobile phase.

Hydrocarbon extraction and analysis was performed as described earlier [25]. Briefly, cells from 10 ml of the culture were collected by centrifugation, and lipids extracted with methanol-chloroform extraction. The chloroform phase was analyzed by gas chromatography-mass spectrometry (GC-MS). The GC-MS analysis was performed with Agilent Technologies 6890N/5975B system, column HP-5MS 30 m x 0.25 mm with 0.25 μm film thickness, He flow 4.7 ml/min, 1μl splitless injection, oven program: 55°C hold 5 min, 55°C - 280°C 20°/min ramp, 280°C hold 3 min. Scan 50-500 m/z, 1.68 scan/sec. Heptadecane peaks were identified and quantified based on external standard (Sigma-Aldrich, USA).

## Declarations

### Authors’ contributions

TL, SS, and VS designed the study. TL performed the molecular, microbiological, and analytical work, analyzed the data, and wrote the manuscript. HV participated in initial strain construction and characterization. All authors read and approved the final manuscript.

### Data availability

The datasets generated and analyzed during the current study are available from the corresponding author on reasonable request.

